# Antimicrobial Peptides Induce Cell Death in Marginal Zone Lymphoma Models Resistant to Targeted Therapies

**DOI:** 10.1101/2025.09.13.674604

**Authors:** Filippo Spriano, Alberto J. Arribas, Fangwen Zhang, Maria Luisa Mangoni, Francesco Buonocore, Francesco Bertoni

**Affiliations:** Institute of Oncology Research (IOR), Faculty of Biomedical Sciences, USI, Bellinzona, Switzerland; Laboratory Affiliated to Pasteur Italia-Fondazione Cenci Bolognetti, Department of Biochemical Sciences, Sapienza University of Rome, Rome, Italy; Department for Innovation in Biological, Agro-Food and Forest Systems, University of Tuscia, Viterbo, Italy; Oncology Institute of Southern Switzerland (IOSI), Ente Ospedaliero Cantonale, Bellinzona, Switzerland

**Keywords:** lymphoma, peptides, resistance, BTK, temporins, trematocines, chionodracines

## Abstract

Marginal zone lymphoma (MZL) is an indolent yet incurable B-cell malignancy in which targeted agents such as BTK and PI3K inhibitors frequently fail due to resistance or toxicity. Antimicrobial peptides (AMPs), evolutionarily conserved effectors of innate immunity, possess selective cytotoxicity against malignant cells by exploiting tumor-specific membrane alterations. We evaluated the antitumor activity of seven natural AMPs, including Antarctic fish-derived trematocines and chionodracine variants, and amphibian temporins, against MZL cell lines (VL51, Karpas1718) and derivatives resistant to BTK, PI3Kδ, or PI3Kα/δ inhibitors. Among them, W-trematocine and temporin L demonstrated potent dose-dependent cytotoxicity with IC_50_ values of 5.7-10 μM, maintaining full activity in all resistant models. Other peptides showed moderate activity, while chionodracine-1 was inactive. Notably, W-trematocine displayed minimal toxicity toward non-malignant cells in prior studies, underscoring its selectivity. AMP-mediated killing, driven by membrane disruption and non-apoptotic death pathways, bypassed conventional resistance mechanisms, suggesting therapeutic potential in relapsed/refractory disease. These findings highlight natural AMPs as promising candidates for development in drug-resistant MZL, warranting further optimization and preclinical validation.

---

Marginal zone lymphoma (MZL) is a heterogeneous group of indolent B-cell non-Hodgkin lymphomas encompassing extranodal mucosa-associated lymphoid tissue (MALT) lymphoma, nodal MZL, and splenic MZL ^1^. Although these lymphomas are generally slow growing, they remain incurable in most patients. In recent years, targeted therapies have substantially improved clinical outcomes. Agents such as Bruton’s tyrosine kinase (BTK) inhibitors (ibrutinib, zanubrutinib), and phosphoinositide 3-kinase (PI3K) inhibitors (idelalisib, copanlisib) have become central to the therapeutic landscape ^1,2^. However, their efficacy is often limited by either the emergence of resistance mutations, activation of alternative signaling pathways, or intolerable toxicities that necessitate treatment discontinuation ^1,2^. Therefore, there is an unmet clinical need to identify novel therapeutic strategies capable of bypassing traditional resistance mechanisms.

Antimicrobial peptides (AMPs) are evolutionarily conserved bioactive molecules that represent one of the most ancient and fundamental strategies of host defence against microbial invasion ^3^. These small peptides (mainly 5 to 50 amino acid residues in their biologically active sequence) are widely distributed across all forms of life, ranging from bacteria and fungi to plants, invertebrates, and vertebrates. Their broader presence highlights a critical role in innate immunity, where they not only function as microbicidal agents but also contribute to immunomodulation, wound healing, and regulation of inflammation. AMPs exert their biological activity primarily by interacting with and disrupting microbial membranes; their amphipathic conformation, coupled with a net positive charge, allows preferential binding to the negatively charged phospholipids present in bacterial cell walls and fungal membranes ^4^. While AMPs have long been studied for their antimicrobial properties, there is now increasing attention toward their ability to induce cytotoxicity in malignant cells, with minimal effects on healthy tissues ^5^. The rationale for this lies in the similarities between microbial and tumor cell membranes. Neoplastic cells, particularly those undergoing rapid proliferation, frequently carry a net negative charge due to exposure of phosphatidylserine, altered glycosylation patterns, and increased heparan sulfate proteoglycans ^6^. These changes provide a biochemical basis for selective targeting of cancer cells by cationic AMPs, while sparing most normal cells. AMPs generally compromise plasma membrane integrity, induce mitochondrial dysfunction, or trigger non-apoptotic forms of cell death such as necrosis ^5,7^. These unique mechanisms raise the possibility that AMPs could be particularly useful in drug-resistant cancers, where canonical apoptotic pathways are impaired.

We investigated the potential of natural AMPs to exert cytotoxic activity against MZL cell models, including variants resistant to currently available targeted agents. Seven peptides of natural origin were selected based on their previously reported antimicrobial or antitumor properties. These included three peptides from Antarctic fish, two variants of trematocine derived from *Trematomus bernacchii*, and one variant of a peptide, chionodracine, isolated from *Chionodraco hamatus*, as well as four temporins from amphibians (*Rana temporaria*) ^8–11^ (Figure 1). The Antarctic peptides are of particular interest because organisms inhabiting extreme environments often evolve structurally unique peptides with enhanced stability and bioactivity ^8–9^. Temporins, in turn, are among the smallest AMPs and have been extensively studied for their amphipathic α-helical conformations, which favor interactions with membranes^7^.

**Figure 1.**
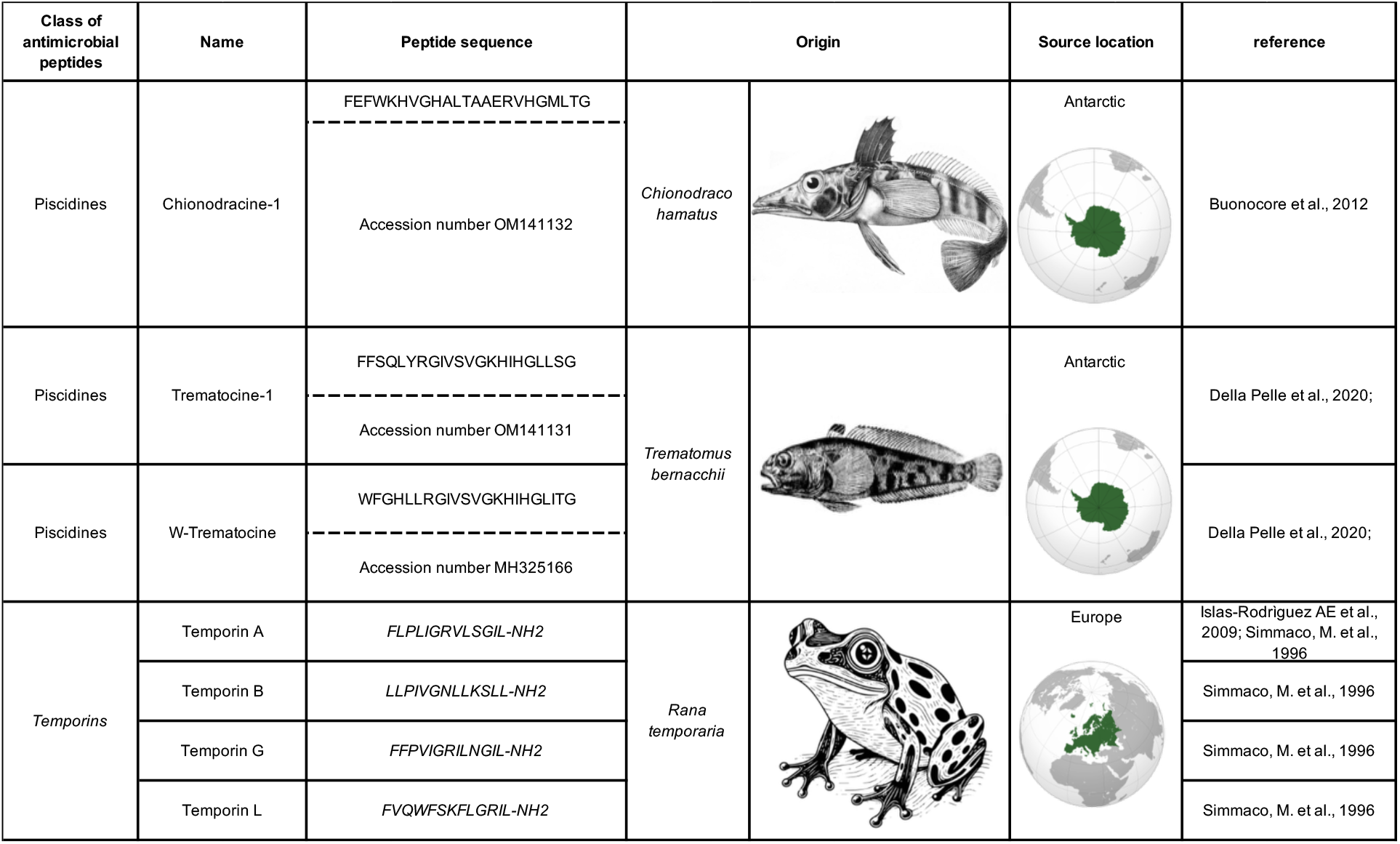
Antimicrobial peptides (AMPs) and species from which the peptides are derived. Peptides derived from *Chionodraco hamatus* (chionodracine peptide), *Trematomus bernacchii* (trematocine peptides), and *Rana temporaria* (temporin peptides) were tested for their potential antitumor activity. The geographic distribution of each species is highlighted on the maps. Created in https://BioRender.com.

For *in vitro* testing, we employed two human MZL cell lines, VL51 and Karpas1718, alongside derivative clones rendered resistant through prolonged exposure to the BTK inhibitor ibrutinib, the PI3Kδ inhibitor idelalisib, or the PI3Kα/δ inhibitor copanlisib (Supplementary Table S1; methods described in Supplementary Materials). Our results revealed a differential spectrum of activity among the tested AMPs. Chionodracine-1 exhibited no measurable cytotoxic effect at concentrations up to 50 µM. In contrast, W-trematocine, a variant of trematocine enriched with a tryptophan residue, and temporin L demonstrated pronounced dose-dependent cytotoxicity in both parental and resistant VL51 and Karpas1718 cells (Figures 2; Supplementary Table S2). The calculated IC_50_ values ranged from 5.7 to 10 µM (Supplementary Table S2). Notably, the cytotoxic effect was fully preserved in all resistant cell line derivatives, suggesting that the mechanism of action of these peptides bypasses the canonical signaling pathways targeted by BTK, PI3K, and BCL2 inhibitors. Additional peptides, including trematocine-1 and other temporin variants, demonstrated moderate cytotoxic activity, with IC_50_ values in the range of 25–50 µM (Figures 2–3), suggesting a potential structure-activity relationship deserving of further optimization. Importantly, earlier work had shown that W-trematocine displays low toxicity toward non-malignant cells. Specifically, exposure of primary human fibroblasts and rabbit erythrocytes to trematocine resulted in negligible cytotoxicity (up to 25μM, 24h) and minimal hemolysis, respectively ^9^. These results underscore the distinct mode of action of AMPs compared with conventional small-molecule inhibitors. While acquired resistance to BTK or PI3K inhibitors often involves secondary mutations in the drug-binding site, activation of alternative kinases, or evasion of apoptosis through upregulation of anti-apoptotic proteins, these adaptations are unlikely to protect tumor cells from AMP-mediated membrane disruption. In addition, the preserved activity of AMPs in resistant cell lines suggests that these peptides could serve as salvage therapies for relapsed/refractory MZL, either as monotherapy or in rational combinations.

**Figure 2.**
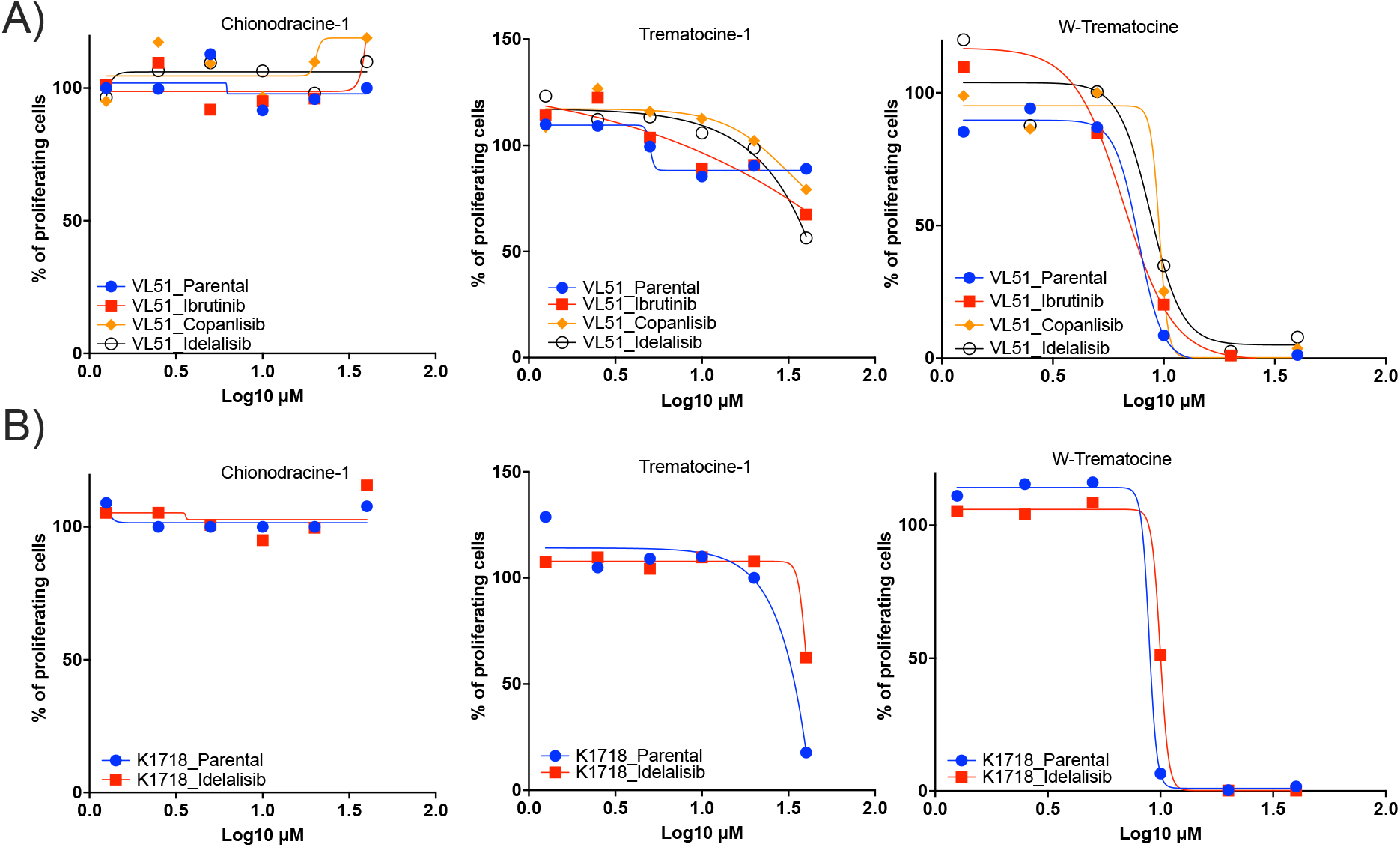
Dose-response curves of chionodracine and trematocine antimicrobial peptides (AMPs) in VL51 and Karpas1718. A) Parental and drug-resistant (idelalisib, ibrutinib, copanlisib) VL51 cells, and B) parental and idelalisib-resistant Karpas1718 cells treated with chionodracine and trematocine derived peptides. Cell viability was assessed after treatment with increasing peptide concentrations. Y-axis: Percentage of proliferating cells relative to the control. X-axis: Compound concentrations (in μM) plotted on a Log10 scale. Methods are described in Supplementary Materials.

**Figure 3.**
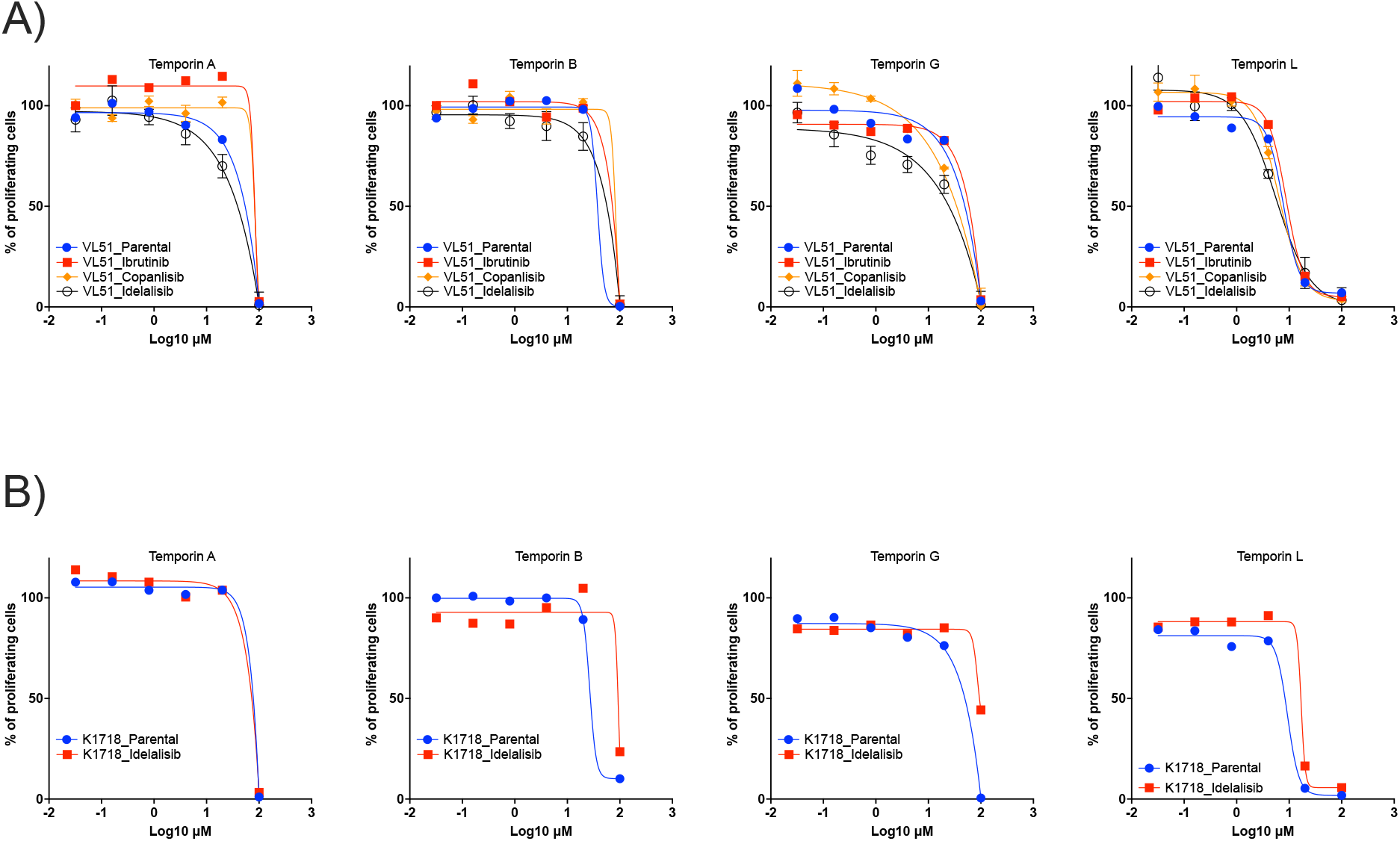
Dose-response curves of temporins antimicrobial peptides (AMPs) in VL51 (A) and Karpas1718 (B). A) Parental and drug-resistant (idelalisib, ibrutinib, copanlisib) VL51 cells, and B) Parental and Idelalisib-resistant Karpas1718 cells treated with temporins. Cell viability was assessed after treatment with increasing peptide concentrations. Data are presented as mean ± standard deviation (error bars). Y-axis: Percentage of proliferating cells relative to the control. X-axis: Compound concentrations (in μM) plotted on a Log10 scale. Methods are described in Supplementary Materials.

In conclusion, our study demonstrates that natural AMPs, particularly W-trematocine and temporin L, exert potent antitumor effects against marginal zone lymphoma cells, including variants resistant to BTK, PI3K, and BCL2 inhibitors. These findings position AMPs as promising candidates for the development of novel therapeutics in drug-refractory B-cell malignancies. Further work is warranted to optimize their pharmacological properties, delineate more precisely their mechanisms of action, and evaluate their efficacy in preclinical lymphoma models. Given their unique mode of action and preserved activity in resistant disease, AMPs may ultimately complement or expand the current therapeutic.

## Funding

This work was partially supported by institutional research funds from the Swiss National Science Foundation (SNSF 31003A_163232/1), Swiss Cancer Research (KFS-4727-02-2019). FZ was supported by the China Scholarship Council (CSC) and the Swiss State Secretariat for Education, Research and Innovation (SERI).

## Author Contributions

FS: performed experiments, interpreted data, and co-wrote the manuscript.

AJA: performed experiments and interpreted data.

FZ: performed experiments.

MLM, FBu: co-designed the study, provided reagents, interpreted data and provided advice.

FB: co-designed the study, interpreted data, supervised the study, and co-wrote the manuscript. All authors reviewed and accepted the final version of the manuscript.

## Conflict of interest

AJA: travel grant from AstraZeneca, consultant fee for PentixaPharm.

FBe: institutional research funds from ADC Therapeutics, Bayer AG, BeiGene, Floratek Pharma, Helsinn, HTG Molecular Diagnostics, Ideogen AG, Idorsia Pharmaceuticals Ltd., Immagene, ImmunoGen, Menarini Ricerche, Nordic Nanovector ASA, Oncternal Therapeutics, Spexis AG; consultancy fee from BIMINI Biotech, Floratek Pharma, Helsinn, Immagene, Menarini, Vrise Therapeutics; advisory board fees to institution from Novartis; expert statements provided to HTG Molecular Diagnostics; travel grants from Amgen, AstraZeneca, iOnctura.

The other Authors have nothing to disclose.

