## Supplementary materials for "Antimicrobial Peptides Induce Cell Death in Marginal Zone Lymphoma Models Resistant to Targeted Therapies"

### Supplementary Methods

Cell lines

The established human cell lines VL51 [1] and Karpas1718 [2], derived from bona fide marginal zone lymphomas (MZL), and their derivatives with secondary resistance obtained by long exposure to the BTK inhibitor ibrutinib, the PI3Kδ inhibitor idelalisib or the PI3Kα/δ inhibitor copanlisib (three from VL51 [3-5], and one from Karpas1718 [6]). Cell lines were cultured in RPMI supplemented with fetal bovine serum (FBS) (10%) and penicillin-streptomycin-neomycin (≈5,000 units penicillin, 5 mg streptomycin, and 10 mg neomycin/mL; Sigma-Aldrich, Darmstadt, Germany). Cell line identities were confirmed by periodically testing them for short tandem repeat (STR) DNA fingerprinting using the Promega GenePrint 10 System kit (B9510). Cells were regularly tested for mycoplasma negativity using the MycoAlert Mycoplasma Detection Kit (Lonza, Visp, Switzerland).

Peptides

Temporins were obtained from Biomatik (Wilmington, DE, USA) and synthesized using solid-phase Fmoc chemistry. Purity (>95%) was confirmed by reverse-phase high-performance liquid chromatography (RP-HPLC) while molecular masses were verified by mass spectrometry. Peptides were dissolved in nuclease-free water, and 2 mM stock solutions were prepared [7].
Antarctic peptides were synthesized by Caslo Aps (Caslo Aps Kongens, Lyngby, Denmark) with a purity of 98%. Purity was confirmed by reverse-phase high-performance liquid chromatography (RP-HPLC) while molecular masses were verified by mass spectrometry. As previously reported, the peptide stock concentration in nuclease-free water (1 mM) was spectrophotometrically determined [8].

*In vitro* cytotoxic activity

Cells were manually seeded into 96-well plates (VL51, 10’000 cells per well; SSK41, 20’000 cells per well; Karpas1718, 30’000 cells per well). Treatments were applied manually, and after 72 hours, cell viability was assessed using 3-(4,5-dimethylthiazol-2-yl)-2,5-diphenyltetrazolium bromide (MTT). Following a 4-hour incubation, the reaction was terminated by adding sodium dodecyl sulfate (SDS) lysis buffer. Plates were analyzed with Cytation3, and IC50s were calculated with the 4-parameter logistic model (R environment). Antarctic peptides were used at a concentration of 40μM, 1:2 dilutions, down to 625 nM. Temporins were used at a concentration of 100μM, 1:5 dilutions, down to 32 nM.

### Supplementary Tables

**Supplementary Table S1. MZL models used in the work.**

| Mode | Exposure to | Sensitivity to BTK inhibitors | Sensitivity to BCL2 inhibitors | Sensitivity to PI3K inhibitors | References |
|---|---|---|---|---|---|
| VL51 | - | yes | yes | yes | [1] |
| Karpas1718 | - | yes | yes | yes | [2] |
| VL51 | Idelalisib | yes | reduced | no | [3] |
| VL51 | Copanlisib | yes | no | no | [4] |
| VL51 | Ibrutinib | no | yes | reduced | [5] |
| Karpas1718 | Idelalisib | no | no | no | [6] |

**Supplementary Table S2. IC50 values obtained by exposing MZL cells for 72 hours to seven antimicrobial peptides**

| | **Chionodracine-1** | **Trematocine-1** | **W-Trematocine** | **Temporin A** | **Temporin B** | **Temporin G** | **Temporin L** |
|---|---|---|---|---|---|---|---|
| **Cell lines** | IC50 (μM) | IC50 (μM) | IC50 (μM) | IC50 (μM) | IC50 (μM) | IC50 (μM) | IC50 (μM) |
| VL51 | >40 | >40 | 6.9 | 32.0 | 38.2 | 49.7 | 7.6 |
| Karpas1718 | >40 | >40 | 8.7 | 47.4 | 44.1 | 38.6 | 5.7 |
| VL51-Ide | >40 | >40 | 9.8 | 27.5 | 31.6 | 20.2 | 6.3 |
| Karpas1718-Ide | >40 | >40 | 10.1 | 49.0 | 71.4 | 80.5 | 9.7 |
| VL51-Copa | >40 | >40 | 8.5 | 48.8 | 47.7 | 35.1 | 7.1 |
| VL51-Ibru | >40 | >40 | 7.4 | 48.5 | 45.4 | 33.9 | 9.5 |
